# The short term High Fat Diet sparks gut microbiota associated metabolic syndrome pathogenesis and impact of Amoxicillin intervention in mice

**DOI:** 10.1101/2021.09.22.461387

**Authors:** Suresh Kumar, Vikram Saini, V. Samuel Raj

## Abstract

**Background:** Knowledge about alteration in gut microbiota on a high-fat diet (HFD) and common antibiotics for a short period is little known. The study was undertaken to evaluate the changes in the microbiota profile in HFD fed mice followed by amoxicillin intervention.

**Results:** For two weeks, the mice were fed with HFD followed by amoxicillin for one week. Animals were evaluated for haemato-biochemical, histopathological and 16S rRNA sequencing followed by bioinformatics analysis to know any changes in gut microbiota ecology. Amoxicillin treatment for a short duration had marked the remarkable remodelling of gut microbiota that directly linked to beneficial impact by countering the pathogenesis of the metabolic syndrome besides overgrowth of particular opportunistic pathogens of gut inflammatory diseases. Amoxicillin treatment significantly decreased blood glucose and a slight elevation in cholesterol levels on blood biochemistry analysis. No marked pathological changes were observed in HFD fed mice.

**Conclusions:** Our results suggest that Amoxicillin treatment had a beneficial influence on the obese related metabolic syndrome with the risk of hypercholesterolemia and some gut-resistant intestinal pathobionts.

## Background

The global problem of consumption of high fat, a portion of westernized food, has serious adverse effects on human metabolic and immune health. High-fat diet (HFD) sparks chronic low-grade systemic inflammation cascading that modulate the gut microbiota by increasing the levels of endotoxins, circulating free fatty acids, and inflammatory mediators that finally impede the homeostasis of many organs [1]. Many studies have explained that prolonged high-fat consumption (8–24 weeks) modulates mice gut microbiota composition and functionality [2]. Evidence is now persuasive that the use of antibiotics could have various adverse outcomes, including microbial dysbiosis, decreased colonization resistance in the gut microbiota and altered metabolic activity such as the swing of biochemical parameters [3]. Nowadays, metabolic syndrome is a severe health condition associated with mainly high blood sugar, abnormal cholesterol or triglyceride levels and chronic inflammation linked to alteration of gut microbiota. Antibiotics target the bacteria that cause infections and alter the resident microbiota, affecting host physiology and potentially host health. Antibiotics medication in metabolic syndromes might increase or decrease the risk by selectively altering these circulating parameters associated with remodelling of gut microbiota. None of the information is available about the short term effect of a high-fat diet on gut microbiota profile and metabolic changes. We somewhat anticipated the short term intake of HFD even induces physiopathology of metabolic syndrome by structural and functional disruption of gut microbiota. However, the impact of antibiotic treatment would be either beneficial or detrimental to the circulating parameters of the host depending upon the change of gut microbiota species. This taking into account, we examined the changes in the caecal content microbiota composition by employing 16S rRNA high-throughput sequencing technology as well as the effect on haematological and biochemical parameters and vital organs including heart, liver and kidney in mice after a short period of HFD feeding (2 weeks) followed by Amoxicillin treatment intervention (1 week).

## Results

### Haematological parameters

Haematological parameters are shown in Figure 1. The significant decrease of haematocrit (HCT), means corpuscular volume (MCV) and white blood cells (WBCs) was observed in the CD vs HFD (*p*=0.022, 0.02189, 0.0128). A significant increase of thrombocytes was observed in the CD vs HFD (*p*=0.0001). Interestingly, the thrombocytes on Amoxicillin treatment significantly decreased (*p*=0.0001); however, it did not substantially alter HCT and MCV as observed in the HDF + Amox group.

**Figure 1.**
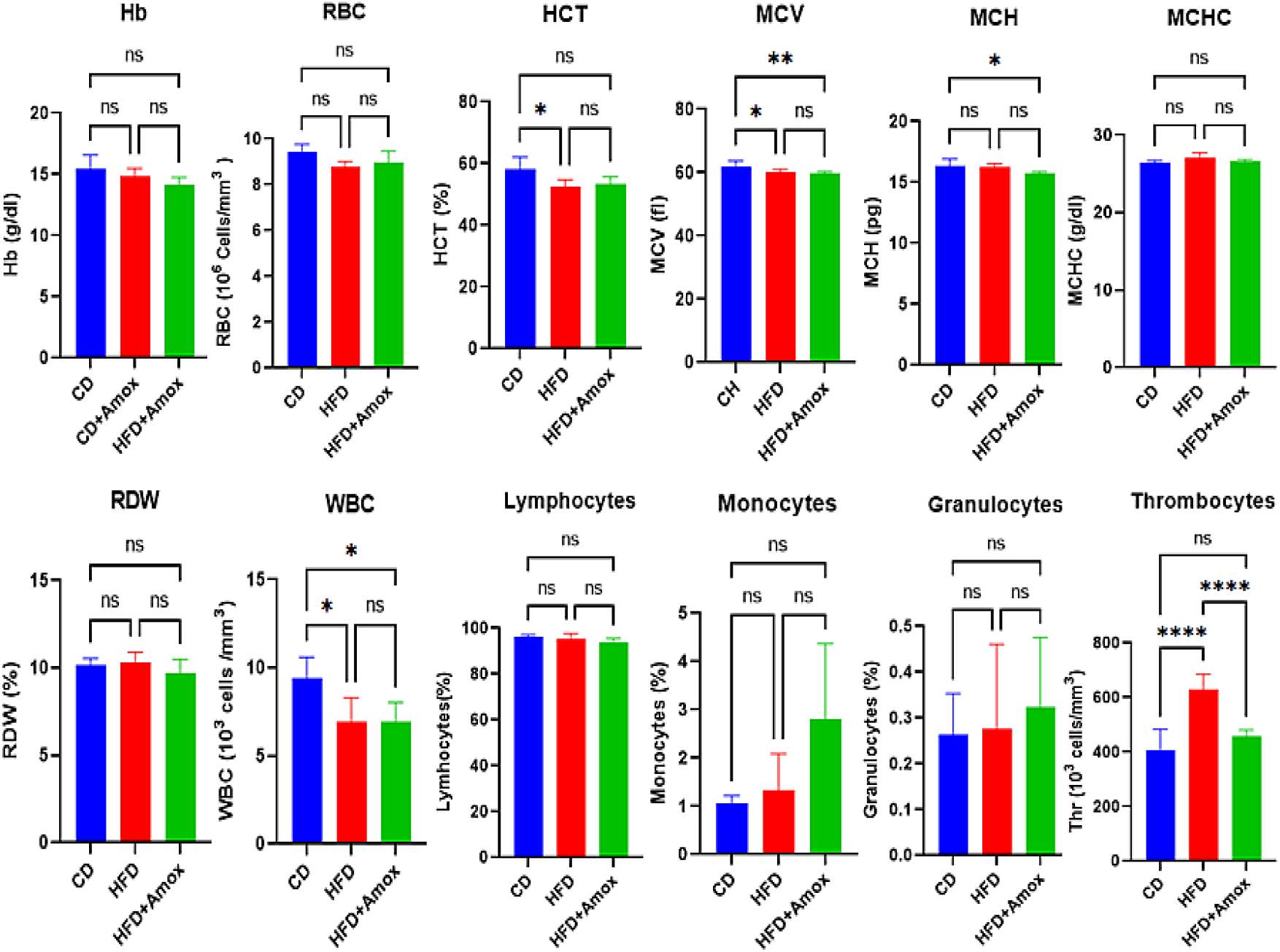
Short term effect of a high-fat (45%) diet followed by Amoxicillin treatment on haematological parameters in mice Values are means ± SD for 6 samples in each group. Statistically significant differences were determined by one way ANOVA followed by Bonferroni test: * p < 0.05, from CD: Standard chow diet; HFD: High-fat diet; HFD+ Amox.: High-fat diet+Amoxicillin

### Biochemical parameters

This study examined critical biochemical parameters in the animals’ serum, such as total cholesterol, total triglycerides (TG), alanine aminotransferase (ALT), aspartate aminotransferase (AST), creatinine, urea, and glucose (Figure 2). No significant changes were observed in ALT, AST, and creatinine concentration in the CD versus HFD. Compared to HFD, the HFD+Amox showed a significantly reduced ALT and AST concentration (p=0.0015, 0.0001). There was also a significant increase in total cholesterol and glucose level in the CD versus HFD (p<0.0001). Amoxicillin treatment significantly reduced glucose levels, as observed in the HFD vs HFD+Amox (*p* <0.0001), but no significant change in cholesterol levels was observed. This study also recorded a substantial reduction in the triglycerides and urea levels in the CD versus HFD (*p*<0.0001). However, Amoxicillin treatment did not significantly change the concentration of triglycerides and urea, as noticed in the HFD vs HFD+Amox group.

**Figure 2.**
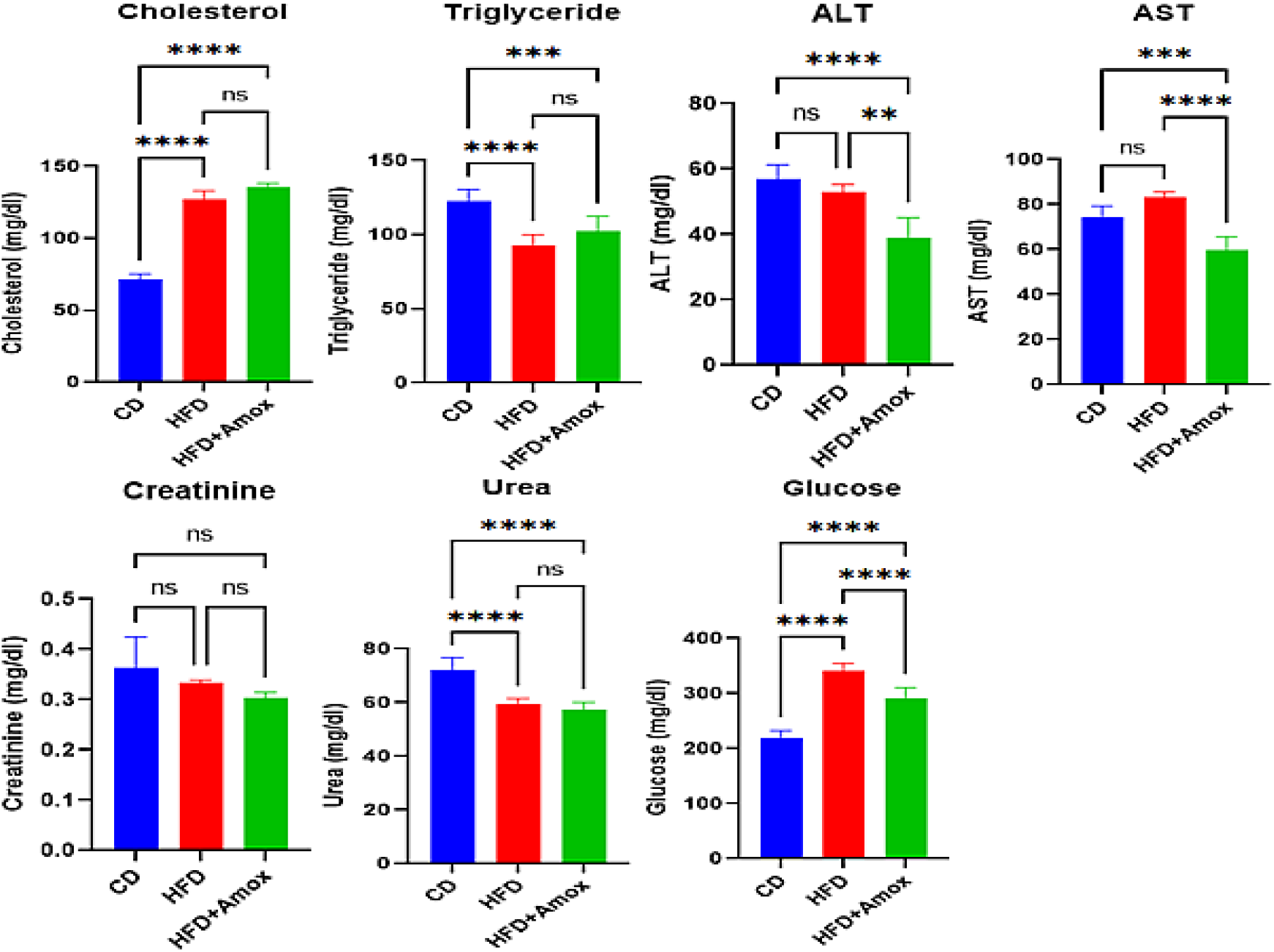
Short term effect of high-fat (45%) diet followed by Amoxicillin treatment on biochemical parameters in mice. Values are means ± SD for 6 samples in each group expressed as mg/dL. Statistically significant differences were determined by one-way ANOVA followed by Bonferroni test: * p < 0.05, from CD: Standard chow diet; HFD: High-fat diet; HFD+ Amox.: High-fat diet+Amoxicillin

### Gut microbiota

After filtering, 878130 high-quality sequences were produced, with an average of 292710 reads per sample. The total number of OTUs at the 97% similarity level was 121183, as shown in Table 1. The Shannon-Wiener curve of all three samples already reached a plateau at this sequencing depth (Figure 3), suggesting that the sequencing was deep enough. Estimators of the alpha and beta diversity are and summarized in Tables 1 and 2 and shown in Figure 3

**Table 1.**
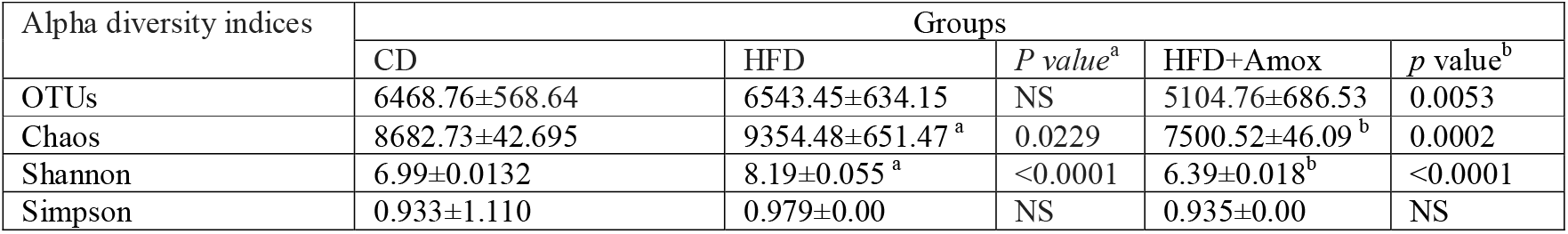
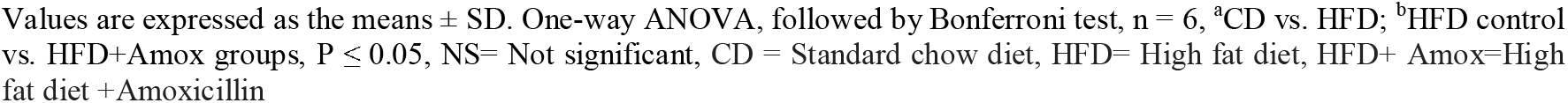
Short term effect of high fat (45%) diet followed by amoxicillin treatment on alpha diversity

| Alpha diversity indices | Groups |  |  |  |  |
| --- | --- | --- | --- | --- | --- |
|  | CD | HFD | *P value*<sup>a</sup> | HFD+Amox | *p value*<sup>b</sup> |
| OTUs | 6468.76±568.64 | 6543.45±634.15 | NS | 5104.76±686.53 | 0.0053 |
| Chaos | 8682.73±42.695 | 9354.48±651.47<sup>a</sup> | 0.0229 | 7500.52±46.09<sup>b</sup> | 0.0002 |
| Shannon | 6.99±0.0132 | 8.19±0.055<sup>a</sup> | <0.0001 | 6.39±0.018<sup>b</sup> | <0.0001 |
| Simpson | 0.933±1.110 | 0.979±0.00 | NS | 0.935±0.00 | NS |

Values are expressed as the means ± SD. One-way ANOVA, followed by Bonferroni test, $n = 6$ , <sup>a</sup>CD vs. HFD; <sup>b</sup>HFD control vs. HFD+Amox groups, $P \leq 0.05$ , NS= Not significant, CD = Standard chow diet, HFD= High fat diet, HFD+ Amox=High fat diet +Amoxicillin

**Table 2.** Short term efect of high fat diet (45%) diet followed by amoxicillin treatment on beta diversity

| Beta diversity indices | Groups |  |
| --- | --- | --- |
|  | CD vs HFD | HFD vs HFD+Amox |
| Unweighted UniFrac distance | 0.618607754 | 0.627751097 |
| Weighted UniFrac distance | 0.23798881 | 0.371335623 |
| Bray-Curtis dissimilarity | 0.70783875 | 0.874340147 |
| Jaccard Abundance | 0.190967872 | 0.222592696 |

CD = Standard chow diet, HFD= High fat diet; HFD+ Amox=High fat diet +Amoxicillin

**Figure 3.**
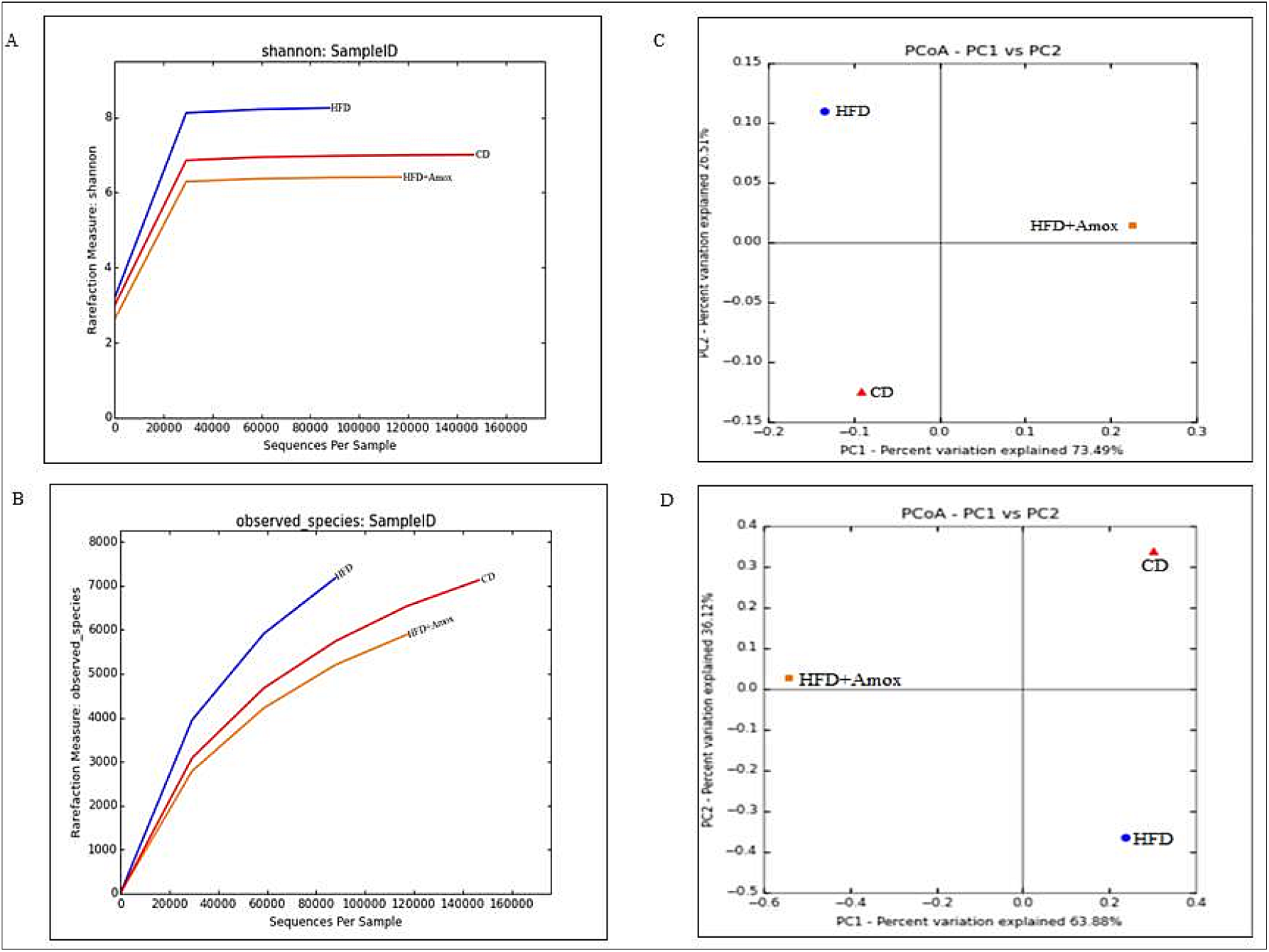
Rarefaction curves and Shannon-Wiener curves of each sample.Shannon-Wiener curves (A) and rarefaction curves (B) were all calculated at the 97% similarity level with pyrosequencing data in microbiota from groups of CD, HFD and HFD+Amox. Principal coordinates analysis (PCoA) based on Weighted UniFrac (C) and Bray-Curtis dissimilarity matrix (D) for three groups. CD = Standard chow diet, HFD= High fat diet, HFD+ Amox=High fat diet +Amoxicillin

### Taxonomy-based comparisons of gut microbiota at the phylum and genus levels among three groups

Overall microbiota compositions for each group at the phylum in presented in Tables 3 and shown in Figure 4. The microbiota compositions at Family levels is are presented in Table 4. *Firmicutes, Proteobacteria, Bacteroides* and *Actinobacteria* were the dominant phylum in these groups. *Firmicutes/ Bacteroidetes* (*F/B* ratio) was significantly higher in the HFD (48.3), with a value more than thrice that of the CD (15.46). In contrast, it significantly decreased more than twice in the HFD+Amox (19.3) compared to the HFD group. Core bacteria analysis revealed that gut dominant phylum *Firmicutes* was significantly more abundant in the CD (80.4%) and HFD+Amox (73.7%), compared to that in the HFD (67.7%). Similarly, the *Bacteroides* were significantly more abundant in CD (5.25 %;) relative to HFD (1.4%) and HFD+Amox (3.8%). In contrast, the relative abundance of *Proteobacteria* was significantly increased in HFD (13.8%), as compared to CD (7.5%) and HFD+Amox (9.2%). Interestingly, *Actinobacteria* were significantly more abundant in HFD (9.6%), compared to CD (3.8%) and HFD+Amox (9.1%).

**Table 3.** Effect of high fat (45%) diet followed by amoxicillin treatment on microbiota at phylum level in mice

| Phylum | Relative abundance (%) |  |  |  |  |
| --- | --- | --- | --- | --- | --- |
|  | CD | HFD | *p* value<sup>a</sup> | HFD+Amox | *p* value<sup>b</sup> |
| *Firmicutes* | 80.4± 0.396 | 67.70± 0.467<sup>a</sup> | <0.0001 | 73.70± 0.440<sup>b</sup> | <0.0001 |
| *Proteobacteria* | 7.50±0.897 | 13.80± 0.344<sup>a</sup> | <0.0001 | 9.20 ±0.289<sup>b</sup> | <0.0001 |
| *Bacteroidetes* | 5.20±0.233 | 1.40±0.117<sup>a</sup> | <0.0001 | 3.80 ±0.191<sup>b</sup> | <0.0001 |
| *Actinobacteria* | 3.80±0.191 | 9.60 ±0.294<sup>a</sup> | <0.0001 | 9.10 ±0.287<sup>b</sup> | 0.0143 |
| *Verrucomicrobia* | 0.20±0.044 | 0.30±0.054 | NS | 0.30±0.054 | NS |
| *TM7* | 0.10±0.031 | 3.30±0.178<sup>a</sup> | <0.0001 | 0.10±0.007<sup>b</sup> | <0.0001 |
| *Acidobacteria* | 0.30±0.054 | 0.60 ±0.077<sup>a</sup> | <0.0001 | 0.60±0.077 | NS |
| *Cyanobacteria* | 0.50±0.070 | 0.30±0.054<sup>a</sup> | 0.0002 | 0.40±0.063<sup>b</sup> | 0.0435 |
| *Firmicutes/Bacteroidetes (F/B) ratio* | 15.46±0.361 | 48.35±0.449<sup>a</sup> | <0.0001 | 19.30±0.394<sup>b</sup> | <0.0001 |

Values are expressed as the means ± SD. One-way ANOVA, followed by Bonferroni test, n = 6, <sup>a</sup>CD vs. HFD; <sup>b</sup>HFD control vs. HFD+Amox groups, P ≤ 0.05, NS= Not significant, CD = Standard chow diet, HFD= High fat diet, HFD+ Amox=High fat diet +Amoxicillin

**Table 4.** Effect of high fat (45%) diet followed by amoxicillin treatment on microbiota at family level in mice

| Phylum | Family | Relative abundance (%) |  |  |  |  |
| --- | --- | --- | --- | --- | --- | --- |
|  |  | CD | HFD | p value<sup>a</sup> | HFD+Amox | p value<sup>b</sup> |
| Firmicutes | Enterococcaceae | 0.00±0.000 | 0.10±0.031 | NS | 3.10±0.173<sup>b</sup> | <0.0001 |
| Firmicutes | Lactobacillaceae | 4.50±0.207 | 15.20±0.359<sup>a</sup> | <0.0001 | 0.30±0.054<sup>b</sup> | <0.0001 |
| Firmicutes | Unc Clostridiales | 39.0±0.487 | 32.60±0.468<sup>a</sup> | <0.0001 | 11.2±0.315<sup>b</sup> | <0.0001 |
| Firmicutes | Clostridiaceae | 5.90±0.235 | 4.90±0.222<sup>a</sup> | <0.0001 | 0.30±0.054<sup>b</sup> | <0.0001 |
| Firmicutes | Lachnospiraceae | 12.40±0.329 | 5.70±0.231<sup>a</sup> | <0.0001 | 17.70±0.381<sup>b</sup> | <0.0001 |
| Firmicutes | Peptostreptococcaceae | 0.10±0.031 | 0.10±0.031 | NS | 4.70±0.211<sup>b</sup> | <0.0001 |
| Firmicutes | Ruminococcaceae | 17.40±0.379 | 6.30±0.242<sup>a</sup> | <0.0001 | 1.40±0.117<sup>b</sup> | <0.0001 |
| Firmicutes | Erysipelotrichaceae | 0.30±0.054 | 0.80±0.089<sup>a</sup> | <0.0215 | 33.50±0.471<sup>b</sup> | <0.0001 |
| Proteobacteria | Desulfovibrionaceae | 3.00±0.170 | 7.50±0.263<sup>a</sup> | <0.0001 | 0.20±0.141<sup>b</sup> | <0.0001 |
| Proteobacteria | Enterobacteriaceae | 0.70±0.083 | 0.90±0.094<sup>a</sup> | 0.0442 | 3.30±0.178<sup>b</sup> | <0.0001 |
| Bacteroidetes | Bacteroidaceae | 0.20±0.044 | 0.20±0.044 | NS | 1.40±0.117<sup>b</sup> | <0.0001 |
| Bacteroidetes | S24-7 | 4.10±0.198 | 0.30±0.054<sup>a</sup> | <0.0001 | 1.40±0.117<sup>b</sup> | <0.0001 |
| Actinobacteria | Coriobacteriaceae | 1.40±0.117 | 6.70±0.250<sup>a</sup> | <0.0001 | 5.70±0.231<sup>b</sup> | <0.0001 |

Values are expressed as the means ± SD. One-way ANOVA, followed by Bonferroni test, n = 6, <sup>a</sup>CD vs. HFD; <sup>b</sup>HFD control vs. HFD+Amox groups, P ≤ 0.05, NS= Not significant, CD = Standard chow diet, HFD= High fat diet, HFD+ Amox=High fat diet +Amoxicillin

**Figure 4.**
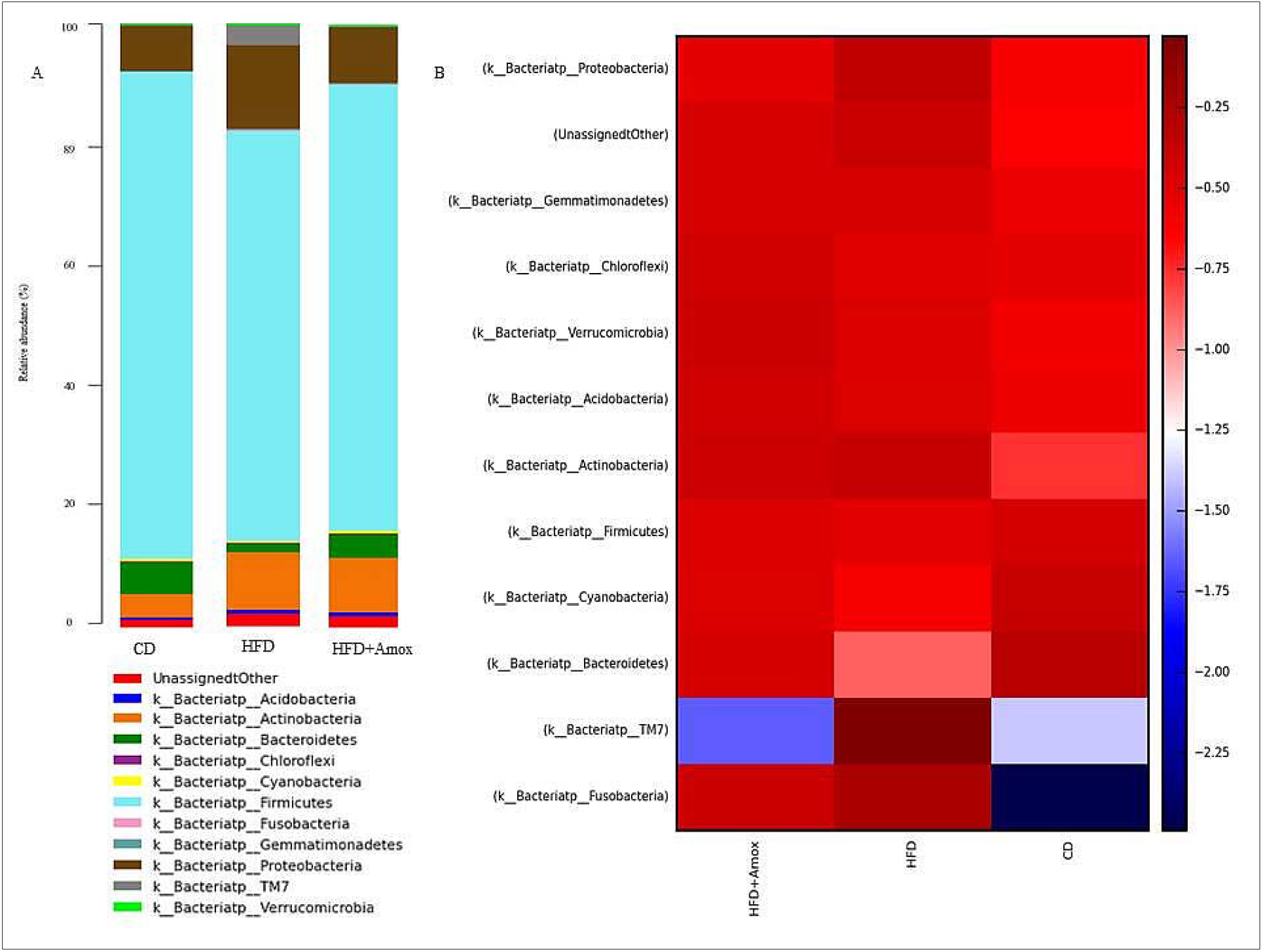
Effect of high fat (45%) diet followed by amoxicillin treatment on microbiota at phylum level in mice CD = Standard chow diet, HFD= High fat diet, HFD+ Amox=High fat diet +Amoxicillin

At the genus level, a total of 21, 40, and 37 genera (Table 5 and Figure 5) were found in CD, HFD, and HFD+Amox group samples, respectively. Compared to CD, HFD fed mice showed considerable increased the relative abundance of *Lactobacillus* (4.50% to 15.20%), *Desulfovibrio* (3.00% to 7.40%), *Oscillospira* (1.80% to 3.10%), *Adlercreutzia* (1.30% to 3.90%), *Rothia* (0.00% to 0.70%) *Streptococcus* (0.00% to 0.60%), *Turicibacter* (0.00% to 0.50%), *Allobaculum* (0.00% to 0.40%) and *Granulicatella* (0.00% to 0.20%). However, Amoxicillin treatment in HFD mice caused considerable decrease of *Lactobacillus* (15.20% to 0.30%), *Desulfovibrio* (7.40% to 0.20%), *Oscillospira (*3.10% to 0.80%), *Rothia (*0.70% to 0.60%) *Streptococcus* (0.60% to 0.50%), *Turicibacter* (0.50% to 0.00%), *Allobaculum* (0.04% to 0.00%), G*ranulicatella* (0.20% to 0.10%) and increased of *Adlercreutzia* (3.90% to 5.60%).

**Table 5.**
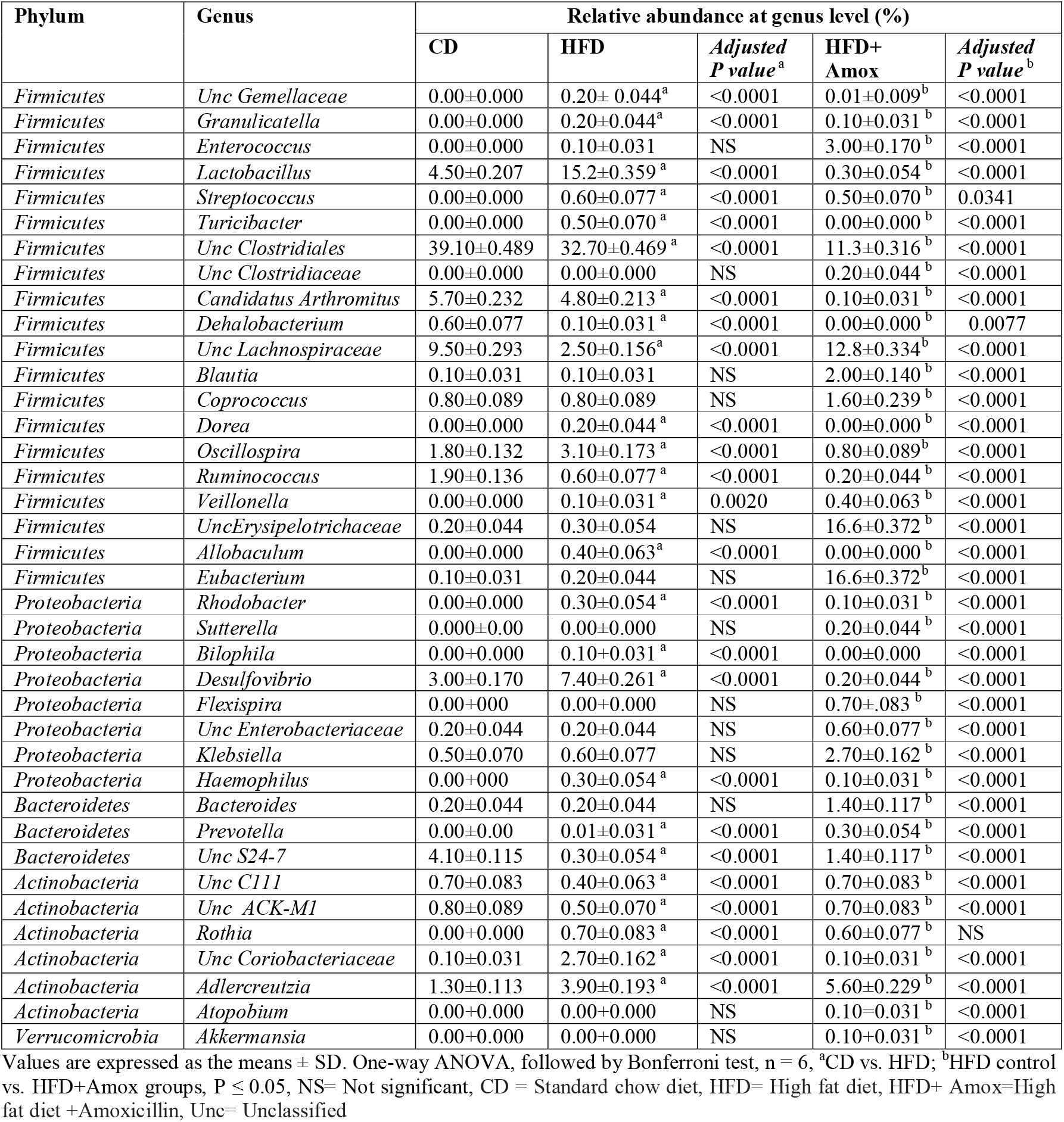
Effect of high fat (45%) diet followed by amoxicillin treatment on microbiota at genus level in mice

**Figure 5.**
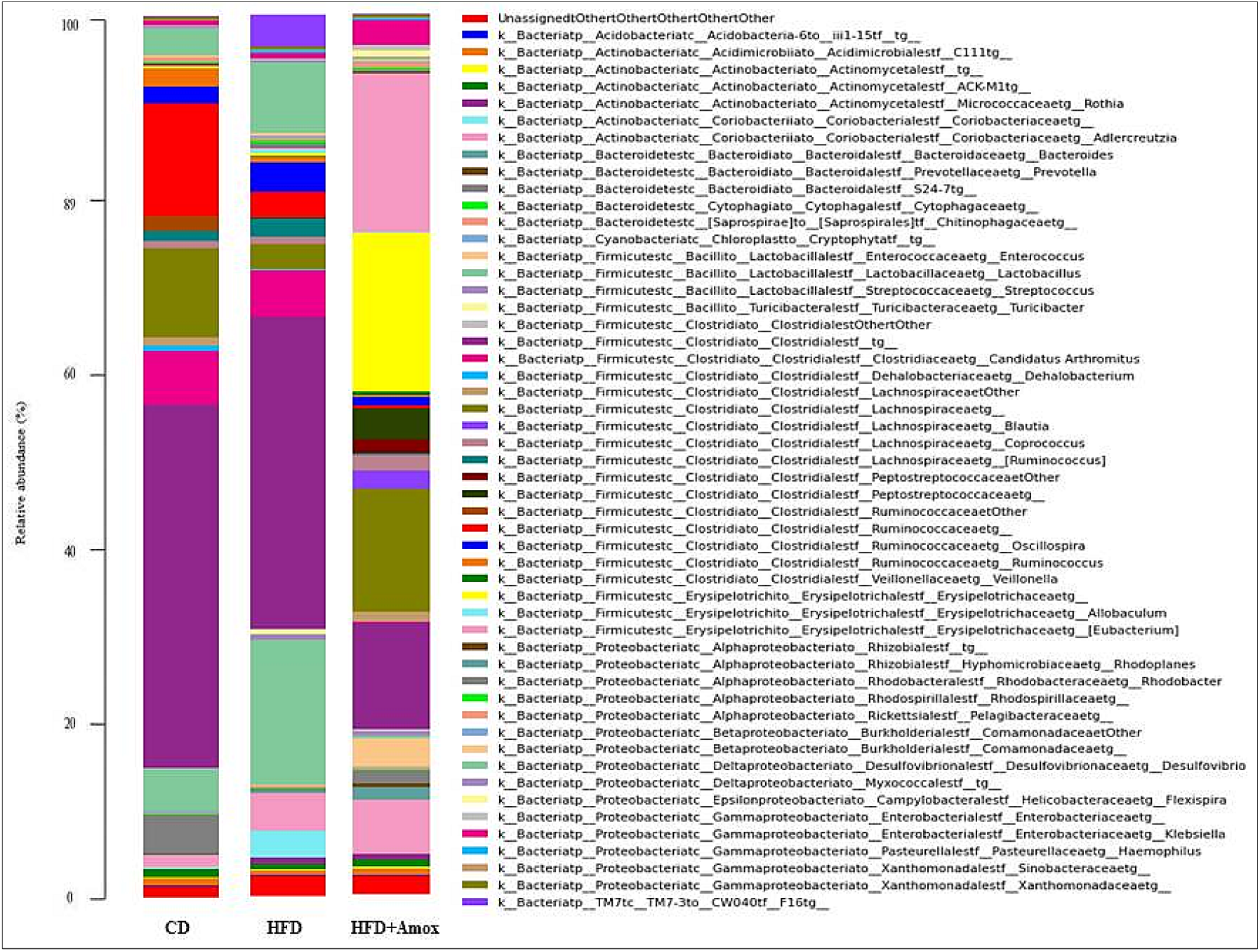
Effect of high fat (45%) diet followed by amoxicillin treatment on microbiota at genus level in mice CD = Standard chow diet, HFD= High fat diet; HFD+ Amox=High fat diet +Amoxicillin

HFD fed mice compared to CD revealed an increase in the abundance of unclassified *Erysipelotrichaceae* (0.10% to 0.20%), *Eubacterium* (0.10% to 0.20%), *Enterococcus* (0.00% to 0.10%), *Veillonella* (0.00% to 0.10%), *Klebsiella* (0.50% to 0.60%) and *Prevotella* (0.00% to 0.10%). Amoxicillin treated HFD mice showed further increase in the abundance of *Erysipelotrichaceae* (0.30% to 16.6%), *Eubacterium* (0.20% to 16.60%), *Enterococcus* (0.10% to 3.00%), *Veillonella* (0.10% to 0.40%), *Klebsiella* (0.60% to 2.70%), and *Prevotella* (0.10% to 0.30%).

HFD fed mice compared to CD showed a decreased in the abundance of *Unclassified Clostridiales* (39.10% to 32.7%), *Unclassified Lachnospiraceae* (9.50% to 2.50%), *Unclassified S24-7* (4.10% to 0.30%), *Ruminococcaceae* (1.90% to 0.60%) and *Candidatus Arthromitus* (5.70% to 4.80%). Amoxicillin treated HFD mice showed an increase in the relative abundance of *Unclassified Lachnospiraceae* (9.50% to 2.50%), *Unclassified S24-7* (4.10% to 0.30%), while decreased of *Ruminococcaceae* (0.60% to 0.20%), and *Candidatus Arthromitus* (4.80% to 0.10%).

Compared to HFD, HFD+Amox showed an increase in *Bacteroides* (0.20% to 1.40%), *Coprococcus* (0.80% to 1.60%), *Flexispira* (0.00% to 0.70%), *Blautia* (0.01% to 2.00%), *Sutterella* (0.00% to 0.20%), *and Unclassified Enterobacteriaceae* (0.02% to 0.70%). However compared with CD, HFD group samples did not show any considerable changes. Further, the important microbiota *Akkermansia muciniphila* (0.01%) was observed in HFD+Amox group only.

The liver of the HFD fed mice (Figure 6C) showed mild congestions in the sinusoidal, central & portal veins. Polyhedral shape hepatocytes with slightly vacuolated granular cytoplasm and vesicular nuclei were also observed. Focal area of mild degenerative changes with swollen hepatocytes & some cells with karyolytic nuclei were also detected. The heart section in HFD demonstrated the congestion in blood vessels with mild hyperplasia in cardiac muscle fibres (Figure 6A). Amoxicillin-treated HFD fed mice showed normal histological architecture of the heart, liver and kidney (Figure 6B, 6D and 6F).

**Figure 6.**
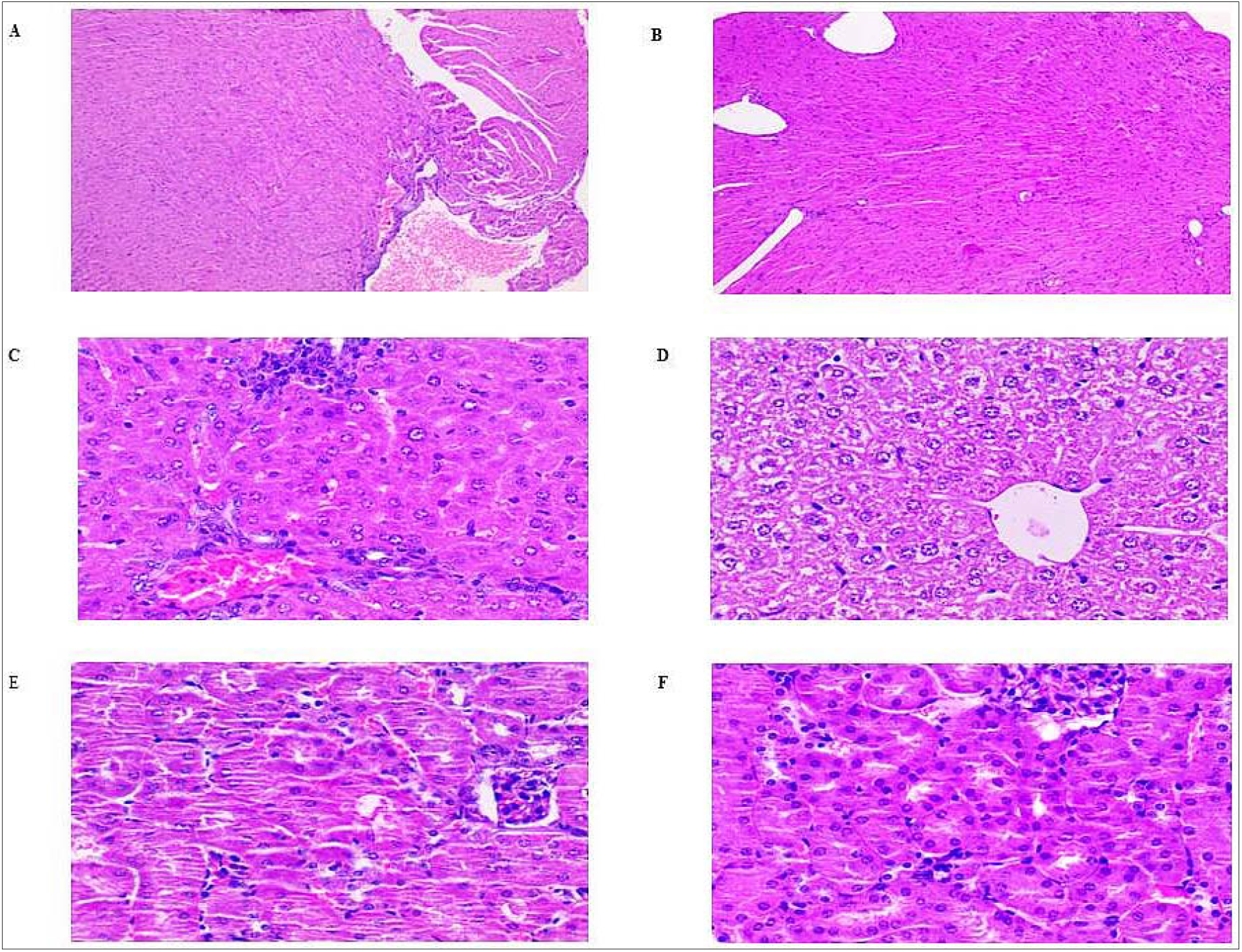
Short term effect of high fat (45%) diet followed by amoxicillin treatment on mice’s heart, liver, and kidney. The heart section of the HFD group demonstrated the congestion in blood vessels with mild hyperplasia in cardiac muscle fibres (A). Liver histology in the HFD group (C) showed mild sinusoidal congestion of the central & portal vein. Notice swollen hepatocytes & some of the cells with karyolytic nuclei. Also, observe focal infiltration of inflammatory cells and micro abscess. The kidney section in the HFD group showed normal architectue (E). The heart, liver and kidney histology in HFD+Amox showed no sign of morphological changes (B, D, & F). CD = Standard chow diet, HFD= High fat diet; HFD+ Amox=High fat diet +Amoxicillin

## Discussion

Metabolic syndrome is a severe health condition that fastens people’s lives at higher risk across the world. Further, the swing of metabolic parameters by antibiotics in metabolic disorders warns the researchers and clinicians. We conducted a short HFD feeding for two weeks followed by an Amoxicillin intervention for one week in mice. Our study has provided additional evidence that HFD for a short period in mice models could substantially change the relative abundance of microbiota composition at the genus and the phylum level [4]. The highest taxonomic richness, evenness and diversity were observed in HFD fed mice, and lowest in Amoxicillin treated HFD fed mice, as indicated by alpha diversity estimators.

This study exhibited a significant increase in *F/B* ratio, associated with the pathology of obese associated metabolic syndrome with a decrease in *Firmicutes* and *Bacteroidetes i*n HFD fed mice [5]. Consistently, this study also observed the significsnt increase in *Proteobacteria* and *Actinobacteria o*n HFD fed mice [5,6]. Amoxicillin treatment in HFD fed mice showed a reverse trend of significantly decreased *F/B* ratio by considerably rising in the abundance of *Firmicutes* and *Bacteroidete*s along with a significant decline in *Proteobacteria* and *Actinobacteria*. Our results suggest that Amoxicillin tenement may be partially beneficial in the inflammatory state of a metabolic syndrome rather than further deteriorating the pathogenesis implicated with most antibiotics.

Our study was congruent with the previous findings of the significant increase in the abundance of *Lactobacillus, Desulfovibrio, Oscillospira, Streptococcus, Granulicatella, Turicibacte*r, *Allobaculum*, and *Rothia* in HFD fed mice [7,8,9,10]. The above gut bacteria played the mechanisms that have a significant role in developing metabolic syndrome. Most of the strains of *Lactobacillus* nourish gut health by secreting antimicrobial substances in response to invaders that stop pathogens from colonizing the gut. The most striking feature in this study HFD fed mice showed the diminution of *Lactobacillus reuteri*, which helps maintain the gut epithelial barrier and prevents the growth of pathobionts in the intestine by secreting antibiotic substances such as reuteri [11,12]. The increase of *Desulfovibrio* develops obesity-related metabolic disorders by subsiding butyrate production and increased Hydrogen sulfide and lipopolysaccharide (LPS). These modulations finally drive insulin resistance and disruption of the immune system [13,14]. The abundance of *Oscillospira* positively correlates with fasting serum insulin and reduction in *Zonula occludens-1 (ZO-1*) mRNA expression indicates their potential role in developing obesity-related metabolic disorders and gut dysfunction [14,15]. *Streptococcus* produces fermentation of sugars and yields lactic acid linked with developing metabolic diseases [16]. The enrichment of *Granulicatella* was associated with obesity [17].

There was a positive correlation between the *Allobaculum* and the expression of *Angiopoietin-related protein (4ANGPTL4*) that plays a crucial role in lipid deposition by repressing the *lipoprotein lipase (LPL)* enzyme and developing obesity-related metabolic disorders [18]. The cholesterol abundance in the HFD diet might be the reason for the increase in *Turicibacter* [19]. *Rothia* is known for its versatile metabolic capacities and provides growth substrates to Gram-negative non-mucus-degrading bacteria, a more specialized community including *Pseudomonas aeruginosa* [20]. In the present study, Amoxicillin treatment in HFD fed mice showed a significant decrease in *Lactobacillus, Desulfovibrio, Oscillospira, Streptococcus, Granulicatella, Turicibacte*, and *Allobaculum*. The relative abundance of Rothia was decreased but not found significantly. *Desulfovibrio* were the most susceptible to Amoxicillin treatment [21. Patrice D *et al*. demonstrated that Antibiotic treatment reduced metabolic endotoxemia resulting in the reduced glucose intolerance, lower inflammation, oxidative stress and macrophages infiltration in visceral adipose tissue [22]. Similarly, Amoxicillin treatment seems to positively impact the metabolic syndrome pathology by improving the insulin signalling and gut epithelium integrity by decreasing the abundance of microbiota linked to obese related metabolic disorders and gut inflammatory diseases.

This study was congruent with the previous results of the significant decrease of *Lachnospiraceae* [23], *S24-7* [2], *Ruminococcaceae* [24] and *Candidatus Arthromitus* [25]. *Ruminococcacea*e and *S 24-7* reduction was found with lower-level butyrate production that induces metabolic endotoxemia and various metabolic disorders. *Lachnospiraceae* produce short-chain fatty acids (SCFA), especially propionate and provide energy for the maintenance of other gut microbes and improve the development of host epithelial cells. Previous gut microbiota studies reported that eradicating *Candidatus arthromitus* in the mouse gut microbiota resulted in metabolic changes leading to obesity. Amoxicillin-treated HFD mice showed the increased relative abundance of members of *Lachnospiraceae* and *S24-7*, while it further decreased the abundance of *Ruminococcacea*e and *Candidatus arthromitus*.

Our study could not detect any alteration in the abundance of *Coprococcus, Blautia Sutterella, Allobaculum*, and *Bacteroides* in HFD fed mice as reported by other studies [7,10,26,27,28]. Interestingly, this study found a significant increase in the abundance of *Coprococcus, Blautia, Sutterell*a, *Bacteroides* and decreased *Allobaculum* on Amoxicillin treated HFD fed mice. The increase in the above bacteria has a beneficial effect as an increase in *Coprococcus*, negatively correlated with hypertension, Basl Metabolic Index (BMI), and body fat percentage [29,30]. *Blautia* abundance was observed with decreased fasting plasma glucose levels, haemoglobin A1C (HbA1c), and inhibited colonization of pathogenic bacteria in the intestine [31,32]. This study detected the increase of *Blautia producta* known for secreting a bacteriostatic substance, lantibiotic, which inhibits the growth of *Vancomycin-resistant enterococci* [VRE] [32]. The actual effect of *Sutterella* on health remains unclear [33]. We also detected the *Bacteroides*, especially *Bacteroides acidifaciens* and. *Bacteroides uniformis* induce GLP-1 activation by using the TGR5 receptor through bile acids, taurine, and cholate that counters obesity and increases insulin sensitivity in mice [34]. *Allobaculum* was decreased in this study as having a positive correlation with expression of *Angiopoietin-relatedprotein (4ANGPTL4)* that cause lipid deposition by repressing the *lipoproteinlipase (LPL)* enzyme and developing obesity-related metabolic disorders [18].Conclusively, Amoxicillin treatment appears to influence metabolic syndrome pathology positively.

Apart from the beneficial effects of shifting gut microbiome by Amoxicillin on HFD fed mice, we also observed an increase in the relative abundance of other gut microbiota such as *Erysipelotrichaceae* [35], *Enterococcus* [9], *Eubacterium* [10], *Klebsiella* [28], and *Prevotella* [9] in HFD fed mice, as well as their further significant increased of abundance in Amoxicillin, treated HFD fed mice also. The rise in these gut microbiota causes metabolic syndrome and intestinal inflammation. The abundance of *Erysipelotrichaceae* strongly correlates with host cholesterol metabolites and developing symptoms of metabolic syndrome [36,37]. *Enterococcus* symbionts with humans can have critical virulence factors that cause gut inflammation by altering bile acid composition [38,39]. Butyrate produced members of *Eubacterium* play a critical role in maintaining a healthy gut. However, gut dysbiosis caused modification of *Eubacterium*, linked with various human disease states [40]. *Klebsiella*, an intestinal pathobiont, observed a rise in inflammatory bowel disease (IBD), necrotizing enterocolitis (NEC) and colorectal cancer (CRC) [41]. *Prevotella* strains also have pathobiont properties that promote diseases like obesity, inflammatory bowel disease, or other inflammatory diseases [42]. It is worth mentioning that we detected *Enterococcus casseliflavus* and *Prevotella melanogenic* enrichment linked with inflammatory diseases [43,44] in Amoxicillin treated HFD mice only. Our data suggest that Amoxicillin treatment in

HFD mice appeared to trigger the proliferation and virulence of these gut-resistant intestinal pathobionts that may even impose an additional risk in metabolic syndrome pathology. Consistently, this study found a significant increase in total cholesterol in HFD fed. Amoxicillin increased the cholesterol level by increasing the hepatic *HMG-CoA reductase* (45, 46]. Our study also showed slight increases in cholesterol levels in Amoxicillin treated HFD fed mice. The increase in the relative abundance of *Erysipelotrichacea*e, as we observed in the Amoxicillin treated HFD fed mice, could be the reason. HFD fed mice also observed a decrease in TG congruence with Gou J *et al*. [47]. The elevated insulin concentrations could inhibit hepatic very low density lipoprotein (VLDL) secretion and stimulate adipose TG uptake as an alternative mechanism for the reduced serum TG levels. Amoxicillin treatment in HFD fed mice did not cause any significant effect on the TG level. HFD fed mice were observed with an increase in AST but not significantly. The accumulation of liver lipids, which could cause damage to cellular homeostasis by causing cytotoxicity, could be the probable reason. Amoxicillin-treated HFD fed mice showed significantly decreased AST and ALT. The reason could be that Amoxicillin might be hepatoprotective in inflammatory conditions.

Consistently, the present study was observed significantly reduced serum urea levels in HFD fed mice [48]. However, long term HFD studies demonstrated the increased serum urea levels that could be due to a more damaging impact on the liver for prolonged exposure [49]. Amoxicillin treated HFD mice did not find a significant effect on urea level. Most of the studies interpreted that long term treatment of HFD increased blood glucose levels [50]. Interestingly, we found that HFD feeding in mice for 14 days increased blood glucose level similar to Louis XL *et al*. reported that short term feeding of HFD up to 17 days also raised glycaemia [51]. It is worth mentioning that Amoxicillin treated HFD mice significantly reduced the glucose level indicating their hypoglycaemic effect. Zarrinpar A *et al*. reported that Amoxicillin causes the depletion of the microbiome, which alters glucose homeostasis by changing colonocyte energy utilization from short-chain fatty acids (SCFAs) to glucose [52]. Similarly, Rodrigues RR *et al*. demonstrated that Antibiotics affect systemic glucose metabolism via shaping gut microbial communities and regulating gene expression programs in the intestine and liver. They explained that improving glucose tolerance by vancomycin can partially be attributed to the increased abundance of A. muciniphila, which is abstaining in some mouse colonies [53]. Based on the finding, we also attempted to correlate the decrease in the glucose level in Amoxicillin treated HFD fed mice. We expect a significant increase in the relative abundance of *Lachnospiraceae, S24-7, Oscillospira* and the exceptional appearance of *Coprococcus, Blautia*, and *Akkermansia muciniphila* in Amoxicillin treated HFD mice only could be the leading cause for the restoration of glucose level towards the normal range. However, this finding has an excellent clinical relevance that Amoxicillin treatment in the metabolic syndrome pathogenesis improves the insulin signalling that might decrease the blood glucose level.

Long-term HFD studies showed a significant decrease in haematological parameters [54,55]. This study observed a slight decline in the value of Hb, RBC, MCH and a significant decrease in HCT and MCV in HFD fed mice. The reason could be that an HFD promotes a low-grade systemic inflammation that elevates inflammatory markers such as *IL-6* and *IL-1*, which provoked *hepcidin* in adipose tissue, decreased iron absorption, and impaired iron fortification effectiveness [56]. Amoxicillin treated HFD mice did not manifest any significant alteration in haematological parameters. Contrary to the previous finding, we observed a significant increase of thrombocytes in mice fed HFD for two weeks. However, Podrini *et al*. demonstrated that prolonged exposure of HFD at least for four weeks requires inducing the depressive effect on thrombocyte counts [57]. The most remarkable finding was that Amoxicillin treatment caused restoration of the thrombocytes towards the CD group by significantly decreasing their value. These conclusions indicated that Amoxicillin might stimulate thrombocytopenia reported in humans through the hapten-dependent antibody process observed in penicillin [58].

A Short-term HFD intake influences the liver’s susceptibility to inflammatory stimuli through the induction of pro coagulation state in the livers of mice [59]. This study also noted the inflammatory damage to the liver and heart in HFD fed mice. Consistently the study did not observe sufficient damage to kidneys [60]. Amoxicillin-treated HFD mice manifest normal architecture of the heart, liver and kidney. These findings show that Amoxicillin treatment shows a hepatoprotective effect in the metabolic syndrome that correlates to the significant decrease of AST and ALT as observed in Amoxicillin treated HFD mice.

In summary, our study found that even short term HFD negatively impact the gut microbiota favoured the pathophysiology of obese related metabolic disorders. Amoxicillin treatment for a short duration showed a positive trend of reshaping microbiota that declines inflammatory state and even swings the metabolic parameters. The limitation of the study was that it had been performed on mice models, and the findings could not be directly applicable to humans. However, this study could be a platform for further research on humans gut microbiota using amoxicillin antibiotics.

## Methods

### Animal model

According to the NIH Guide for the Care and Use of Laboratory Animals (National Research Council), all the experimental procedures were conducted and approved by the Institutional Animal Ethics Committee (IAEC) of All India Institute of Medical Sciences (AIIMS), New Delhi. All mice were maintained at constant temperature (22 ± 3°C) and humidity (40-60%) on a 12 h light/dark cycle with free access to drinking water and rodent diet throughout the study. From the same cohort, twenty-four male mice C57BL/6J mice of 6 weeks old (initial weight 16–18 g) were allocated into two groups (N=12 for each group) as standard chow diet (CD) (10 % energy from fat) and high-fat diet (HFD), HFD (73 % energy from fat) for two weeks. Hereafter, the diet groups were assigned as CD for control and HFD treated mice. Subsequently, half of the animals (N=6) from two groups were euthanized at the end of 2 weeks. The rest half of the animals allocated to each group were fed on their respective diet for a further period of one week. For one week, these animals were treated with amoxicillin at a 50 mg/kg body weight dose, i.e. 0.25 mg/ml in drinking water. After one week of antibiotics, the animals were sacrificed, and samples were collected to see the effect of Amoxicillin on HFD fed mice. The treatment/diet group were referred to as HFD for control and HFD+Amox for treated mice.

### Blood and tissue and caecal samples collection

After two weeks, three mice fasted overnight, and the blood samples were collected via cardiac puncture as a terminal procedure. Following blood collection, the organs (heart, liver and kidney) were isolated for histopathological examination. Cecal content samples were taken shortly after dissection and immediately frozen in liquid nitrogen and stored at -80 °C until further use.

### Haematology and Serum biochemistry

Serum biochemistry was performed using serum auto-analyzer Screen Master 3000, Tulip, Alto Santa Cruz, India, then quantifying Coral GPO-PAP kit (CORAL Clinical systems, Goa, India) using manufacturers’ instructions. The haematology analysis was carried out using an automated vet haematology counter (Melet Schloesing Laboratories, Guwahati, India) as per the manufacturer’s instruction.

### Histopathology

After euthanasia, the liver, kidney and heart tissues were dissected and were fixed overnight in 10% neutral buffer formalin and embedded in paraffin blocks. For haematoxylin-eosin (H & E) staining, tissue sections of 4-5 μm thickness were collected on poly-L-lysine (Sigma) coated slides using a microtome and stained according to standard protocols. The digital images were taken by light microscopy (Olympus CX-29: Olympus Optical Co. Ltd, Tokyo, Japan) and camera (Magnus DC 10).

### 16S rRNA Gene Sequencing

Caecal content samples were sent for next-generation sequencing and genus analysis (DNA Xperts Private Limited). According to the manufacturer’s instructions, the total genomic DNA of gut microbiota was extracted using Qiagen DNA Stool Mini Kit. DNA concentration and integrity were assessed by fluorometer and agarose gel electrophoresis, respectively. The 16S rRNA gene amplicon sequencing was performed on the Illumina MiSeq according to protocols described by [61]. The hypervariable region V4 of the bacterial 16S rRNA gene was amplified by using the primers of the universal primer 515f, 50-GTGCCAGCMGCCGCGGTAA-30; and 806r, 50-GGACTACHVGGGTWTCTAAT-30 on the Illumina MiSeq platform.

### Microbial Bioinformatics Analysis

The raw data were filtered to capture clean reads by eliminating the adapter pollution and low-quality sequences [62]. By using FLASH (Fast Length Adjustment of Short reads v 1.2.1), the high-quality paired-end reads were combined with tags with an average read length of 252 bp [63]. According to their unique barcode and primer, sequences were assigned to each sample using Mothur software (V1.35.1, http://www.mothur.org) and removing the barcodes as well as primers to get the effective clean Tags. The tags were then clustered as OTU (Operational Taxonomic Unit) by scripts of USEARCH (v 7.0.1090) software with a 97% similarity threshold [64].

Taxonomic classification of the representative OTU was assigned by using three commonly used databases (Greengenes, SILVA and Ribosomal Database Project (RDP). Finally, an OTU table and a phylogenetic tree were generated for diversity analysis. Alpha-diversity was calculated by using Chao1, observed species, Shannon indices and Simpson indices. Beta-diversity was measured by calculating the unweighted, weighted UniFrac distances, Bray-Curtis dissimilarity and Jaccard distance. Principal coordinate analysis (PCoA) distance matrices created by QIIME [65].

### Statistical Analysis

Graphpad Prism, Version 9.2.0 (3.2.0) software was used for the statistical analysis of experimental data. Values are expressed as the means ± Standard Deviation (SD). Groups were compared by one-way analysis of variance (ANOVA) followed by the Bonferroni test with a value of p ≤ 0.05 as the cut-off for statistical significance.

## List of abbreviations

CD: Standard chow diet
HFD: High-fat diet
HFD+Amox: High-fat diet+Amoxicillin
HCT: Haematocrit
MCV: Means corpuscular volume
WBC: White blood cells
ALT: Aspartate aminotransferase
AST: Aspartate aminotransferase
TG: Triglycerides
LPS: Lipopolysaccharide
ZO-1: Zonula occludens-1
ANGPTL4: Angiopoietin-related protein
SCFA: Short-chain fatty acids
HbA1c: Haemoglobin A1C
TGR5 receptor: G-protein-coupled bile acid receptor 5
ANGPTL4: Angiopoietin-relatedprotein 4
IBD: Inflammatory bowel disease
NEC: Necrotizing enterocolitis
CRC: Colorectal cancer
VLDL: Very low density lipoprotein
OTUs: Operating taxonomic units
PCoA: Principal coordinate analysis

## Acknowledgements

We greatly appreciate Dr Perumal Nagarajan and Dr. Surender Yadav (National Institute of Immunology, New Delhi, India) for providing experimental and statistical supports.

## Author information

V. Samuel Raj & Vikram Saini contributed equally to this work and are co–first authors of this article

## Contributions

SK conducted the research and analyzed the data. VSR and VS contributed to the conceptualization and design of the research and critically revising the manuscript. All authors read and approved the final manuscript.

## Ethics approval

All care, maintenance, and experimental protocols were approved by Institutional Animal Ethics Committee (IAEC) of All India Institute of Medical Sciences

## Consent for publication

Not applicable

## References

1. Tilg H, Zmora N, Adolph TE, Elinav E. The intestinal microbiota fuelling metabolic inflammation. Nat Rev Immunol. 2020; doi: 10.1038/s41577-019-0198-4.

2. Shang Y, Khafipour E, Derakhshani H, Sarna LK, Woo CW, Siow YL O K. Short Term High Fat Diet Induces Obesity-Enhancing Changes in Mouse Gut Microbiota That are Partially Reversed by Cessation of the High Fat Diet. Lipids. 2017; doi: 10.1007/s11745-017-4253-2.

3. Ramirez J, Guarner F, Bustos Fernandez L, Maruy A, Sdepanian VL, Cohen H. Antibiotics as Major Disruptors of Gut Microbiota. Front Cell Infect Microbiol.2020; doi: 10.3389/fcimb.2020.572912.

4. Wu GD, Chen J, Hoffmann C, Bittinger K, Chen YY, Keilbaugh SA, Bewtra M, Knights D, Walters WA, Knight R, Sinha R, Gilroy E, Gupta K, Baldassano R, Nessel L, Li H, Bushman FD, Lewis JD. Linking long-term dietary patterns with gut microbial enterotypes. Science. 2011; doi: 10.1126/science.1208344.

5. Yin X, Liao W, Li Q, Zhang H, Liu Z, Zheng X, Zheng L, Fen, X. Interactions between resveratrol and gut microbiota affect the development of hepatic steatosis: A fecal microbiota transplantation study in high-fat diet mice. Journal of Functional Foods. 2020; 67:103883.

6. Kim SJ, Kim SE, Kim AR, Kang S, Park MY, Sung MK. Dietary fat intake and age modulate the composition of the gut microbiota and colonic inflammation in C57BL/6J mice. BMC Microbiol. 2019; doi: 10.1186/s12866-019-1557-9.

7. Liu S, Qin P, Wang J. High-Fat Diet Alters the Intestinal Microbiota in Streptozotocin-Induced Type 2 Diabetic Mice. Microorganisms. 2019;doi: 10.3390/microorganisms7060176.

8. Lin H, An Y, Hao F. et al. Correlations of Fecal Metabonomic and Microbiomic Changes Induced by High-fat Diet in the Pre-Obesity State. 2016;doi:10.1038/srep21618.

9. Heisel T, Montassier E, Johnson A, Al-Ghalith G, Lin YW, Wei LN, Knights D, Gale CA. High-Fat Diet Changes Fungal Microbiomes and Interkingdom Relationships in the Murine Gut. mSphere. 2017;doi: 10.1128/mSphere.00351-17.

10. Deshpande NG, Saxena J, Pesaresi TG, Carrell CD, Ashby GB, Liao MK, Freeman LR. High fat diet alters gut microbiota but not spatial working memory in early middle-aged Sprague Dawley rats. PLoS One. 2019; doi: 10.1371/journal.pone.0217553.

11. Yi H, Wang L, Xiong Y, Wen X, Wang Z, Yang X, Gao K, Jiang Z. Effects of barrier function in weaned pigs. J Anim Sci. 2018; doi: 10.1093/jas/sky129.

12. Sun J, Qiao Y, Qi C, Jiang W, Xiao H, Shi Y, Le GW. High-fat-diet-induced obesity is associated with decreased antiinflammatory Lactobacillus reuteri sensitive to oxidative stress in mouse Peyer’s patches. Nutrition. 2016; doi: 10.1016/j.nut.2015.08.020.

13. Nagao-Kitamoto H, Kamada N. Host-microbial Cross-talk in Inflammatory Bowel Disease. Immune Netw. 2017; doi: 10.4110/in.2017.17.1.1.

14. Lam YY, Ha CW, Campbell CR, Mitchell AJ, Dinudom A, Oscarsson J, Cook DI, Hunt NH, Caterson ID, Holmes AJ, Storlien LH. Increased gut permeability and microbiota change associate with mesenteric fat inflammation and metabolic dysfunction in diet-induced obese mice. PLoS One. 2012; doi: 10.1371/journal.pone.0034233.

15. Zhang Q, Xiao X, Li M, Yu M, Ping F, Zheng J, Wang T, Wang X. Vildagliptin increases butyrate-producing bacteria in the gut of diabetic rats. PLoS One. 2017;doi: 10.1371/journal.pone.0184735.

16. He C, Cheng D, Peng C, Li Y, Zhu Y, Lu N. High-Fat Diet Induces Dysbiosis of Gastric Microbiota Prior to Gut Microbiota in Association With Metabolic Disorders in Mice. Front Microbiol. 2018;. doi: 10.3389/fmicb.2018.00639.

17. Yang Y, Cai Q, Zheng W, Steinwandel M, Blot WJ, Shu XO, Long J. Oral microbiome and obesity in a large study of low-income and African-American populations. J Oral Microbiol. 2019; doi: 10.1080/20002297.2019.1650597.

18. Fernández-Hernando C, Suárez Y. ANGPTL4: a multifunctional protein involved in metabolism and vascular homeostasis. Curr Opin Hematol. 2020; doi: 10.1097/MOH.0000000000000580.

19. Dimova LG, Zlatkov N, Verkade H.J, Uhlin BE, Tietge U. High-cholesterol diet does not alter gut microbiota composition in mice. Nutrition & metabolism, 2017; doi:10.1186/s12986-017-0170-x.

20. Uranga CC, Arroyo P Jr, Duggan BM, Gerwick WH, Edlund A. Commensal Oral Rothia mucilaginosa Produces Enterobactin, a Metal-Chelating Siderophore. mSystems. 2020; doi: 10.1128/mSystems.00161-20.

21. Lozniewski A, Labia R, Haristoy X, Mory F. Antimicrobial susceptibilities of clinical Desulfovibrio isolates. Antimicrob Agents Chemother. 2001; doi: 10.1128/AAC.45.10.2933-2935.2001

22. Patrice D. C., Rodrigo B., Claude K., et al. (2008). Changes in gut microbiota control metabolic endotoxemia-induced inflammation in high-fat diet–induced obesity and diabetes in mice. Diabetes, 57 (6) 1470–1481. https://doi.org/10.2337/db07-1403

23. Serino M, Luche E, Gres S, Baylac A, Bergé M, Cenac C, Waget A, Klopp P, Iacovoni J, Klopp C, Mariette J, Bouchez O, Lluch J, Ouarné F, Monsan P, Valet P, Roques C, Amar J, Bouloumié A, Théodorou V, Burcelin R. Metabolic adaptation to a high-fat diet is associated with a change in the gut microbiota. Gut. 2012; doi: 10.1136/gutjnl-2011-301012.

24. Daniel H, Gholami AM, Berry D, Desmarchelier C, Hahne H, Loh G, Mondot S, Lepage P, Rothballer M, Walker A, Böhm C, Wenning M, Wagner M, Blaut M, Schmitt-Kopplin P, Kuster B, Haller D, Clavel T. High-fat diet alters gut microbiota physiology in mice. ISME J. 2014; doi: 10.1038/ismej.2013.155.

25. Tomas J, Mulet C, Saffarian A, Cavin JB, Ducroc R, Regnault B, Kun Tan C, Duszka K, Burcelin R, Wahli W, Sansonetti PJ, Pédron T. High-fat diet modifies the PPAR-γ pathway leading to disruption of microbial and physiological ecosystem in murine small intestine. Proc Natl Acad Sci U S A. 2016; doi: 10.1073/pnas.1612559113.

26. Jang LG, Choi G, Kim SW, Kim BY, Lee S, Park H. The combination of sport and sport-specific diet is associated with characteristics of gut microbiota: an observational study. J Int Soc Sports Nutr. 2019 May 3;16(1):21. doi: 10.1186/s12970-019-0290-y.

27. Zheng Z, Lyu W, Ren Y, Li X, Zhao S, Yang H, Xiao Y. Allobaculum Involves in the Modulation of Intestinal ANGPTLT4 Expression in Mice Treated by High-Fat Diet. Front Nutr. 2021; doi: 10.3389/fnut.2021.690138.

28. Singh RP, Halaka DA, Hayouka Z, Tirosh O. High-Fat Diet Induced Alteration of Mice Microbiota and the Functional Ability to Utilize Fructooligosaccharide for Ethanol Production. Front Cell Infect Microbiol. 2020; doi: 10.3389/fcimb.2020.00376.

29. Kim S, Goel R, Kumar A, Qi Y, Lobaton G, Hosaka K, Mohammed M, Handberg EM, Richards EM, Pepine CJ, Raizada MK. Imbalance of gut microbiome and intestinal epithelial barrier dysfunction in patients with high blood pressure. Clin Sci (Lond). 2018;doi: 10.1042/CS20180087.

30. Naderpoor N, Mousa A, Gomez-Arango LF, Barrett HL, Dekker Nitert M, de Courten B. Faecal Microbiota Are Related to Insulin Sensitivity and Secretion in Overweight or Obese Adults. J Clin Med. 2019; doi: 10.3390/jcm8040452.

31. Inoue., Ohue-Kitano R, Tsukahara T, Tanaka M, Masuda S, Inoue T, Yamakage H, Kusakabe T, Hasegawa K, Shimatsu A, Satoh-Asahara N. Prediction of functional profiles of gut microbiota from 16S rRNA type 2 diabetic patients. Journal of clinical biochemistry and nutrition. 2017;doi:10.3164/jcbn.17-44.

32. Liu X, Mao B, Gu J, Wu J, Cui S, Wang G, Zhao J, Zhang H, Chen W. Blautia-a new functional genus with potential probiotic properties? Gut Microbes. 2021;doi: 10.1080/19490976.2021.1875796.

33. Hiippala K, Kainulainen V, Kalliomäki M, Arkkila P, Satokari R. Mucosal Prevalence and Interactions with the Epithelium Indicate Commensalism of Sutterella spp. Front Microbiol. 2016; doi: 10.3389/fmicb.2016.01706.

34. Yang JY, Lee YS, Kim Y, Lee SH, Ryu S, Fukuda S, Hase K, Yang CS, Lim HS, Kim MS, Kim HM, Ahn SH, Kwon BE, Ko HJ, Kweon MN. Gut commensal Bacteroides acidifaciens prevents obesity and improves insulin sensitivity in mice. Mucosal Immunol. 2017; doi: 10.1038/mi.2016.42.

35. Fleissner CK, Huebel N, Abd El-Bary MM, Loh G, Klaus S, Blaut M. Absence of intestinal microbiota does not protect mice from diet-induced obesity. Br J Nutr. 2010;doi: 10.1017/S0007114510001303.

36. Martínez I, Perdicaro DJ, Brown AW, Hammons S, Carden TJ, Carr TP, Eskridge KM, Walter J. Diet-induced alterations of host cholesterol metabolism are likely to affect the gut microbiota composition in hamsters. Appl Environ Microbiol. 2013; doi: 10.1128/AEM.03046-12.

37. Woting A, Pfeiffer N, Loh G, Klaus S, Blaut M. Clostridium ramosum promotes high-fat diet-induced obesity in gnotobiotic mouse models. mBio. 2014; doi: 10.1128/mBio.01530-14.

38. Sánchez B, Cobo A, Hidalgo M, Martínez-Rodríguez AM, Prieto I, Gálvez A, Martínez-Cañamero M. Influence of the Type of Diet on the Incidence of Pathogenic Factors and Antibiotic Resistance in Enterococci Isolated from Faeces in Mice. Int J Mol Sci. 2019; doi: 10.3390/ijms20174290.

39. Seishima J, Iida N, Kitamura K et al. Gut-derived Enterococcus faecium from ulcerative colitis patients promotes colitis in a genetically susceptible mouse host Genome Biol. 2019;doi.org/10.1186/s13059-019-1879-9.

40. Mukherjee A, Lordan C, Ross RP, Cotter PD. Gut microbes from the phylogenetically diverse genus Eubacterium and their various contributions to gut health. Gut Microbes. 2020; doi: 10.1080/19490976.2020.1802866.

41. Pope JL, Yang Y, Newsome RC, Sun W, Sun X, Ukhanova M, Neu J, Issa JP, Mai V, Jobin C. Microbial Colonization Coordinates the Pathogenesis of a Klebsiella pneumoniae Infant Isolate. Sci Rep. 2019; doi: 10.1038/s41598-019-39887-8.

42. Precup G, Vodnar DC. Gut Prevotella as a possible biomarker of diet and its eubiotic versus dysbiotic roles: a comprehensive literature review. Br J Nutr. 2019; doi: 10.1017/S0007114519000680.

43. Klein G. Taxonomy, ecology and antibiotic resistance of enterococci from food and the gastro-intestinal tract. Int J Food Microbiol. 2003; doi: 10.1016/s0168-1605(03)00175-2.

44. Marietta EV, Murray JA, Luckey DH, Jeraldo PR, Lamba A, Patel R, Luthra HS, Mangalam A, Taneja V. Suppression of Inflammatory Arthritis by Human Gut-Derived Prevotella histicola in Humanized Mice. Arthritis Rheumatol. 2016; doi: 10.1002/art.39785.

45. Kim MH, Lee EJ, Cheon JM, Nam KJ, Oh TH, Kim KS. Antioxidant and hepatoprotective effects of fermented red ginseng against high fat diet-induced hyperlipidemia in rats. Laboratory animal research, 2016; doi:10.5625/lar.2016.32.4.217.

46. Rotimi SO, Ojo DA, Talabi OA, Ugbaja RN, Balogun EA, Ademuyiwa O. Amoxillin-and pefloxacin-induced cholesterogenesis and phospholipidosis in rat tissues. Lipids Health Dis. 2015; doi: 10.1186/s12944-015-0011-8.

47. Guo J, Jou W, Gavrilova O, Hall KD. Persistent diet-induced obesity in male C57BL/6 mice resulting from temporary obesigenic diets. PLoS One. 2009; doi: 10.1371/journal.pone.0005370.

48. Dos Santos Lacerda D, Garbin de Almeida M, Teixeira C, de Jesus A, da Silva Pereira Júnior É, Martins Bock P, Pegas Henriques J et al. Biochemical and Physiological Parameters in Rats Fed with High-Fat Diet: The Protective Effect of Chronic Treatment with Purple Grape Juice (Bordo Variety). Beverages. 2018;doi:10.3390/beverages4040100.

49. Ben Gara A, Ben Abdallah Kolsi R, Chaaben R, Hammami N, Kammoun M, Paolo Patti F, El Feki A, Fki L, Belghith H, Belghith K. Inhibition of key digestive enzymes related to hyperlipidemia and protection of liver-kidney functions by Cystoseira crinita sulphated polysaccharide in high-fat diet-fed rats. Biomed Pharmacother. 2017; doi: 10.1016/j.biopha.2016.11.059.

50. Agardh CD, Ahrén B. Switching from high-fat to low-fat diet normalizes glucose metabolism and improves glucose-stimulated insulin secretion and insulin sensitivity but not body weight in C57BL/6J mice. Pancreas. 2012; doi:10.1097/MPA.0b013e3182243107.

51. Louis XL, Thandapilly SJ, MohanKumar SK, Yu L, Taylor CG, Zahradka P, Netticadan T. Treatment with low-dose resveratrol reverses cardiac impairment in obese prone but not in obese resistant rats. The Journal of nutritional biochemistry. 2012; doi: 10.1016/j.jnutbio.2011.06.010.

52. Zarrinpar A, Chaix A, Xu ZZ, Chang MW, Marotz CA, Saghatelian A, Knight R, Panda S. Antibiotic-induced microbiome depletion alters metabolic homeostasis by affecting gut signaling and colonic metabolism. Nat Commun. 2018; doi: 10.1038/s41467-018-05336-9.

53. Rodrigues RR, Greer RL, Dong X, DSouza KN, Gurung M, Wu JY, Morgun A, Shulzhenko N. Antibiotic-Induced Alterations in Gut Microbiota Are Associated with Changes in Glucose Metabolism in Healthy Mice. Front Microbiol. 2017; doi: 10.3389/fmicb.2017.02306.

54. Rojas JM, Bolze F, Thorup I, Nowak J, Dalsgaard CM, Skydsgaard M, Berthelsen LO, Keane KA, Søeborg H, Sjögren I, Jensen JT, Fels JJ, Offenberg HK, Andersen LW, Dalgaard M. The Effect of Diet-induced Obesity on Toxicological Parameters in the Polygenic Sprague-Dawley Rat Model. Toxicol Pathol. 2018; doi: 10.1177/0192623318803557.

55. Mkandla Z, Mutize T, Dludla PV, Nkambule BB. Impaired Glucose Tolerance is Associated with Enhanced Platelet-Monocyte Aggregation in Short-Term High-Fat Diet-Fed Mice. Nutrients. 2019; doi:10.3390/nu11112695.

56. Aigner ., Feldman A, Datz C. Obesity as an emerging risk factor for iron deficiency. Nutrients. 2014; doi; 10.3390/nu6093587.

57. Podrini C, Cambridge EL, Lelliott CJ, Carragher DM, Estabel J, Gerdin AK, Karp NA, Scudamore CL; Sanger Mouse Genetics Project, Ramirez-Solis R, White JK. High-fat feeding rapidly induces obesity and lipid derangements in C57BL/6N mice. Mamm Genome. 2013; doi: 10.1007/s00335-013-9456-0.

58. Mansour H, Saad A, Azar M, Khoueiry P. Amoxicillin/Clavulanic Acid-induced thrombocytopenia. Hospital pharmacy. 2014;doi:10.1310/hpj4910-956.

59. Nanizawa E, Tamaki Y, Sono R, Miyashita R, Hayashi Y, Kanbe A, Ito H, Ishikawa T. Short-term high-fat diet intake leads to exacerbation of concanavalin A-induced liver injury through the induction of procoagulation state. Biochem Biophys Rep. 2020;doi: 10.1016/j.bbrep.2020.100736.

60. Crinigan Calhoun M, Sweaze, KL. Short-Term High Fat Intake Does Not Significantly Alter Markers of Renal Function or Inflammation in Young Male Sprague-Dawley Rats. Journal of nutrition and metabolism, 2015;doi:10.1155/2015/157520.

61. Caporaso J, Lauber C, Walters W et al. Ultra-high-throughput microbial community analysis on the Illumina HiSeq and MiSeq platforms. ISME J. 2012; doi:10.1038/ismej.2012.8.

62. Fadrosh DW, Ma B, Gajer P et al. An improved dual-indexing approach for multiplexed 16S rRNA gene sequencing on the Illumina MiSeq platform. Microbiome, 2014; doi: 10.1186/2049-2618-2-6.

63. Magoc T, Salzberg SL FLASH: fast length adjustment of short reads to improve genome assemblies. Bioinformatics. 2011; doi: 10.1093/bioinformatics/btr507

64. Edgar RC. UPARSE: highly accurate OTU sequences from microbial amplicon reads. Nature methods, 2013; doi: 10.1038/nmeth.2604.

65. Caporaso J G, Kuczynski J, Stombaugh J, Bittinger K, Bushman FD, Costello EK, Fierer N, Peña AG, Goodrich JK, Gordon JI, Huttley GA, Kelley S T, Knights D, Koenig J E, Ley RE, Lozupone CA, McDonald D, Muegge BD, Pirrung M, Reeder J, Knight R. QIIME allows analysis of high-throughput community sequencing data. Nature methods, 2010;doi:10.1038/nmeth.f.303.

